# Two operational modes of atomic force microscopy reveal similar mechanical properties for homologous regions of dystrophin and utrophin

**DOI:** 10.1101/2024.05.18.593686

**Authors:** Cailong Hua, Rebecca A. Slick, Joseph Vavra, Joseph M. Muretta, James M. Ervasti, Murti V. Salapaka

## Abstract

Duchenne muscular dystrophy (DMD) is a lethal muscle disease caused by the absence of the protein dystrophin. Dystrophin is hypothesized to work as a molecular shock absorber that limits myofiber membrane damage when undergoing reversible unfolding upon muscle stretching and contraction. Utrophin is a dystrophin homologue that is under investigation as a protein replacement therapy for DMD. However, it remains uncertain whether utrophin can mechanically substitute for dystrophin. Here, we compared the mechanical properties of homologous utrophin and dystrophin fragments encoding the N terminus through spectrin repeat 3 (UtrN-R3, DysN-R3) using two operational modes of atomic force microscopy (AFM), constant speed and constant force. Our comprehensive data, including the statistics of force magnitude at which the folded domains unfold in constant speed mode and the time of unfolding statistics in constant force mode, show consistent results. We recover parameters of the energy landscape of the domains and conducted Monte Carlo simulations which corroborate the conclusions drawn from experimental data. Our results confirm that UtrN-R3 expressed in bacteria exhibits significantly lower mechanical stiffness compared to insect UtrN-R3, while the mechanical stiffness of the homologous region of dystrophin (DysN-R3) is intermediate between bacterial and insect UtrN-R3, showing greater similarity to bacterial UtrN-R3.

**Significance:** Duchenne muscular dystrophy (DMD) is a severe muscle wasting disorder caused by mutations in DMD gene encoding dystrophin. Utrophin, a fetal homologue of dystrophin, is under active investigation as a dystrophin replacement therapy for DMD. However, it is still unknown if it can substitute dystrophin mechanically. Here, we report mechanical properties of both utrophin and dystrophin fragments encoding the N terminus through spectrin repeat 3 (UtrN-R3, DysN-R3) using atomic force microscope (AFM) through two operational modes, constant speed and constant force. Our data, consistent across both modes, confirm that UtrN-R3 expressed in bacteria exhibits significantly lower mechanical stiffness than insect UtrN-R3. Additionally, bacterial DysN-R3 lies between bacterial and insect UtrN-R3 in terms of mechanical properties, leaning closer to bacterial UtrN-R3.

## Introduction

Duchenne muscular dystrophy (DMD) is a lethal muscle disease that affects 1:5000 boys born in the United States (1) characterized by progressive muscle degeneration and weakness due to loss of the protein dystrophin (2). Dystrophin is a 427 kDa protein expressed primarily at the muscle cell membrane, or sarcolemma, in striated muscle tissue and is a crucial component of the dystrophin-glycoprotein complex (DGC) (3). It is hypothesized that dystrophin acts as a molecular shock absorber to stabilize the sarcolemma and protect muscle against mechanical forces during muscle stretching and contraction (4–7). Thus, dystrophin deficiency increases sarcolemmal fragility ultimately leading to myofiber rupture and death (8).

Structurally, dystrophin is composed of four major domains: an amino terminal (NT) actin-binding domain (ABD1), a large central rod domain with 24 triple helical spectrin-like repeats (SLRs) interspersed with 4 hinge domains, including a second actin-binding domain (ABD2), a cysteine-rich domain binding with the transmembrane dystroglycan complex, and a carboxy-terminal (CT) domain (Fig. 1a). Overexpression of a naturally homologous protein, utrophin, has been proposed as a potential DMD therapy with the goal of substituting utrophin for dystrophin at the sarcolemma (9).

**Fig. 1.**
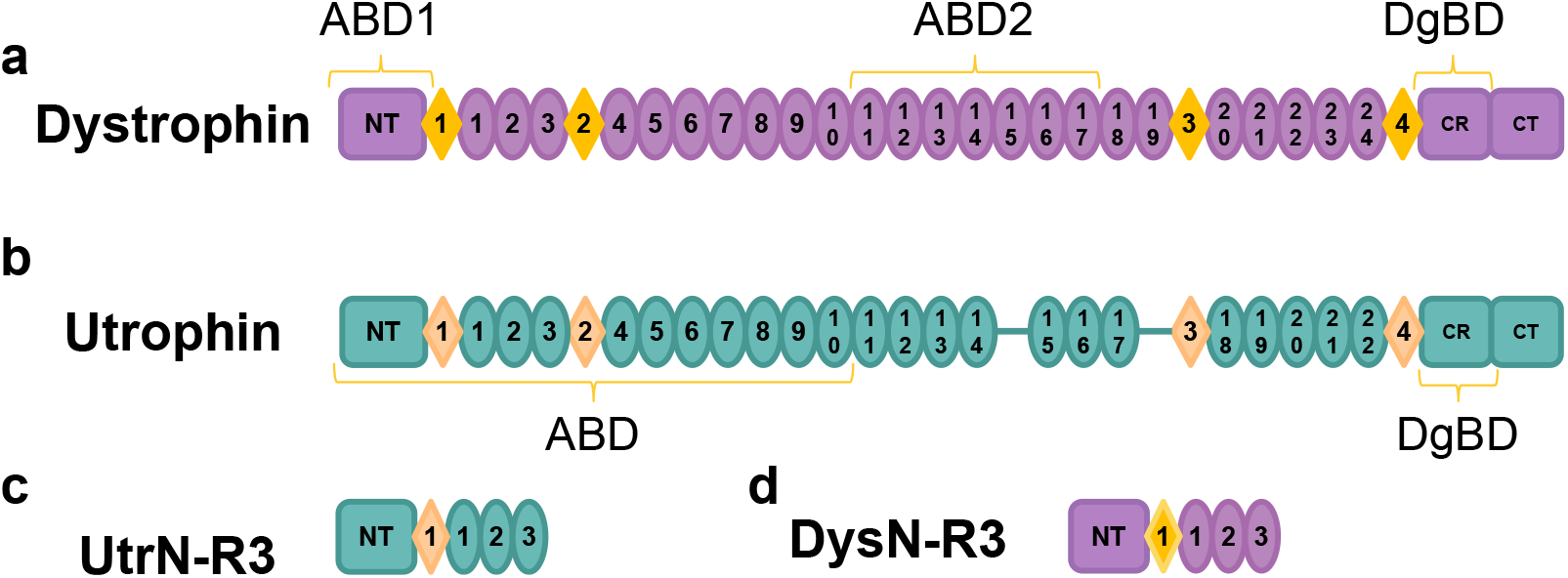
Diagrams of dystrophin and utrophin. (a) Full-length dystrophin. (b) Full-length utrophin. (c) UtrN-R3 construct (d) DysN-R3 construct. Circles, spectrin-like repeats (SLRs); diamonds, hinge domains; NT, N terminus; CT, C terminus; CR, cysteine-rich domain; ABD1 & 2, actin-binding domains; DgBD, dystroglycan binding domain. Figure courtesy of our previous study (13)

Utrophin (Fig. 1b) is abundantly expressed during fetal development, and like dystrophin, forms a complex similar to the DGC at the sarcolemma but is later replaced by dystrophin after birth (10). In adult skeletal muscle, utrophin primarily localizes to the myotendinous and neuromuscular junctions (11). Utrophin overexpression has previously been shown to compensate for dystrophin deficiency in mice (12). While many biochemical investigations of dystrophin and utrophin have been reported, mechanical studies are limited. Therefore, it is yet to be elucidated if utrophin can act as a sufficient mechanical replacement for dystrophin.

Here, we compare the mechanical properties of dystrophin and utrophin fragments encoding the NT through SLR 3 (DysN-R3, Fig. 1c and UtrN-R3, Fig. 1d). We perform single molecule force spectroscopy (SFMS) using constant speed and constant force modes of atomic force microscopy (AFM). In constant speed mode, the molecule is stretched at uniform velocity and the force of domain unfolding is recorded. In constant force mode, a uniform force is imposed on the molecule and the time of unfolding is recorded. We use the constant speed and constant force data to estimate the energy landscapes of the domains, which both cross validate the data from the two modes of operation and provide a framework for understanding the unfolding process (14, 15). The Dudko Hummer Szabo (DHS) model (16) relates the statistics of force magnitude data from constant speed mode to the time to unravel statistics of the constant force mode, retaining applicability regardless of the specific shape of the energy landscape. Our data show consistent results using constant speed and constant force modes. The conclusions drawn from these experimental data are further corroborated by Monte Carlo simulations conducted after recovering energy landscape parameters of the domains.

Our data demonstrate that the mechanical properties of UtrNR3 expressed in bacteria are significantly less mechanically stiff compared to insect UtrN-R3, which is consistent with our previous findings (13). However, our previous study only reported data at one specific speed. In this study, we validate results using force magnitude data from constant speed mode for a range of regulated speeds, the time to unravel statistics from constant force mode for a range of regulated forces, and subsequently recovered energy landscapes. Furthermore, our data on bacterial DysN-R3 suggest that its mechanical properties are significantly lower than those of insect UtrN-R3 but exhibit some similarity to bacterial UtrN-R3.

## Materials and Methods

### Cloning

Mouse utrophin N-R3 (UtrN-R3) was cloned as previously described (13). Similarly, a full-length mouse dystrophin construct was utilized to clone the mouse dystrophin N-R3 (DysN-R3) truncation vector containing an N-terminal FLAG-tag for affinity purification (17, 18).

### Protein expression and purification

Sf9 cells used for insect derived proteins were maintained at 1 × 10^6^ cells/mL in Sf-900 TM II SFM supplemented with penicillin/streptomycin (Sigma-Aldrich) and fungizone (Thermo Fisher Scientific) in a 250 mL suspension culture. For protein expression, 250 mL 1×10^6^ cells/mL cultures were transfected with 10 mL of conditioned media baculovirus and incubated at 150 rpm for 72 hours at 27°C. BL21 E. coli containing N-R3 expression plasmids were grown in selective media for bacterial derived protein expression. Cultures were grown until an *A*_600_ of 0.6-0.8 was reached and protein over-expression was induced by incubating with 1 mM isopropyl-*β*-d-1-thiogalactopyranoside (IPTG, Roche) for 16 hours at 18°C.

Insect and bacterial cells were harvested by centrifuging at 4000 × g for 10 minutes at 4°C. Insect cell pellets were resuspended in lysis buffer containing protease inhibitors and lysed by rotating for 1 hour at 4°C as previously described (13). Lysis buffer was used to resuspend bacterial cell pellets which were lysed by sonication using an ultrasonic homogenizer (BioLogics) with 9 × 1 minute bursts at 40% power and 50% pulse. Each lysate was then centrifuged at 20,000 × g for 10 minutes (insect) or 25 minutes (bacteria) at 4°C to clear the lysates. Cleared supernatant was applied to an anti-FLAG M2 agarose column (Sigma Aldrich) and further purified, dialyzed, and concentrated as described in (13). Protein concentration was determined by Bradford assay using an Albumin standard curve. Purified N-R3 proteins were run on a 4-20% sodium dodecyl sulfate (SDS) polyacrylamide gradient gel and stained with Coomassie blue stain for verification (Fig. S1). Coomassie stained gels were visualized and imaged using Licor Odyssey® Infrared Imaging System.

### Atomic force microscopy experiments

Single molecule force experiments were conducted using MFP-3D atomic force microscope (AFM) (Asylum Research, an Oxford Instruments Company, Santa Barbara, CA). The AFM configuration consists of a flexible cantilever with a sharp tip, a laser-photodiode based sensor for tracking the cantilever tip’s position, and a piezoelectric nano-positioner capable of moving the substrate in three dimensions relative to the cantilever base (19). We employed two types of cantilevers; Bruker MLCT-BIO triangular cantilevers, made of silicon nitride and coated with reflective gold on the backside, with a nominal spring constant of 10 *pN/nm* and a typical tip radius of 20 *nm*, and Olympus BL-RC-150VB cantilevers with Cr/Au coating on the tip with a typical spring constant of 6 *pN/nm*.

Prior to each experiment, the spring constant was determined by analyzing the thermal reaction of the cantilever deflection (20). A droplet (∼ 100 *μl*) of purified protein suspended in PBS, with a dilution ranging from 30 to 100 nMol, was placed on a freshly cleaved and ionized mica substrate. This setup was incubated for 15 minutes before the actual experiment, allowing the protein to adhere to the mica surface. Subsequently, the droplet containing the protein solution was removed, where the mica substrate was washed with 100 *μl* of PBS to eliminate proteins that had not adhered. A new droplet containing 100 *μl* of PBS was introduced to establish the environment for the force spectroscopy experiments. Repeated approach-retraction cycles were conducted at room temperature, where the cantilever’s tip was pressed against the substrate for a duration of 2 seconds, applying an indentation force of approximately 600 *pN*. Subsequently, in the constant speed mode, the cantilever base was retracted at a speed ranging from 500 to 5000 *nm/s*. Since the pulling force on the molecule is balanced by the force experienced by the cantilever, the force, *F* on the protein was determined as,

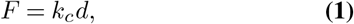

where *k*_*c*_ is the spring constant of the cantilever and *d* is the deflection of the cantilever. In constant force mode, the cantilever base was regulated to maintain a steady deflection, resulting in a regulated force on the molecule, ranging from 40 to 100 *pN*. Data from 300 − 2000 successful force spectroscopy experiments (with at least one identifiable unfolding event) were collected for each protein under a specific speed or force condition and used to determine the unfolding statistics.

### Data analysis

The raw data obtained from the force spectroscopy experiments are primarily composed of three main variables: 1) the time for a bond to unravel and how the forces vary with time, 2) the deflection of the cantilever, and 3) the distance between the base of the cantilever and the substrate. It includes instances where no protein adhered between the tip and substrate, as well as instances where a protein was effectively adsorbed and subsequently stretched. To identify successful experiments and extract meaningful information from data, we utilize a custom algorithm developed in MATLAB R2023b (MathWorks). The algorithm calculates the applied force by multiplying the cantilever deflection with the spring constant and computes the protein extension by subtracting the deflection of the cantilever from the distance between the cantilever base and the substrate. The algorithm first computes the standard deviation *σ* of noise in the force signal measured from the approach curve. Subsequently, in constant speed mode, it detects characteristic peaks in the force signal that exceed 3*σ*. In constant force mode, it identifies steps larger than 10*σ* in the extension signal coinciding with a spike in the force signal. Here, the significant event is the peak in constant speed mode and step in constant force mode. The first significant event arises from the adhesive force between the cantilever tip and the substrate; this event is excluded from the dataset for subsequent analysis. Similarly, the last significant event is often influenced by a detachment event and is discarded. The remaining significant events are recognized as unfolding events of interest. The number of significant events is then used to distinguish between successful and unsuccessful experiments.

The force-extension curves obtained from the constant speed mode are fitted to a worm-like chain (WLC) model. The WLC model (21) relates the force applied on protein *F*_*W LC*_ to its extension *q*, and is given by,

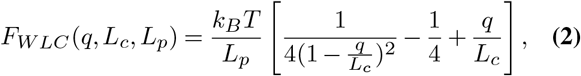

where *k*_*B*_ is the Boltzmann constant, *T* is temperature, and *L*_*c*_ and *L*_*p*_ are the contour length and the persistence length of the protein, respectively. On the other side, extension-time curves obtained from constant force mode are fitted to a staircase function. These curves are then inspected manually. Curves that show irregularities, such as an excessive number of unfolding events or distorted significant events that deviate from the characteristics of the fitting curve, are excluded for further analysis.

### Two state system

In many cases, the unfolding process of a single folded domain of protein can be modeled with a one-dimensional projected energy landscape along the co-ordinate of interest with two stable states being the local minimas in the projected energy landscape. The intrinsic free energy landscape *U*_0_(*x*) along reaction coordinate *x* (see Fig. 2), is assumed to have minimas at folded state *x*_*f*_ and unfolded state *x*_*u*_ with a barrier of height Δ*G*^‡^ at the transition point *x*_*t*_, where Δ*G*^‡^ := *U*_0_(*x*_*t*_) − *U*_0_(*x*_*f*_) is the energy difference between energy levels at *x*_*t*_ and *x*_*f*_. The distance between the folded state and the transition point is defined as Δ*x*^‡^ := *x*_*t*_ − *x*_*f*_. Additionally, the rate of transition from the folded to the unfolded state under intrinsic energy landscape *k*_0_ is exponentially related to the energy barrier, where 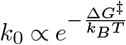 (22).

**Fig. 2.**
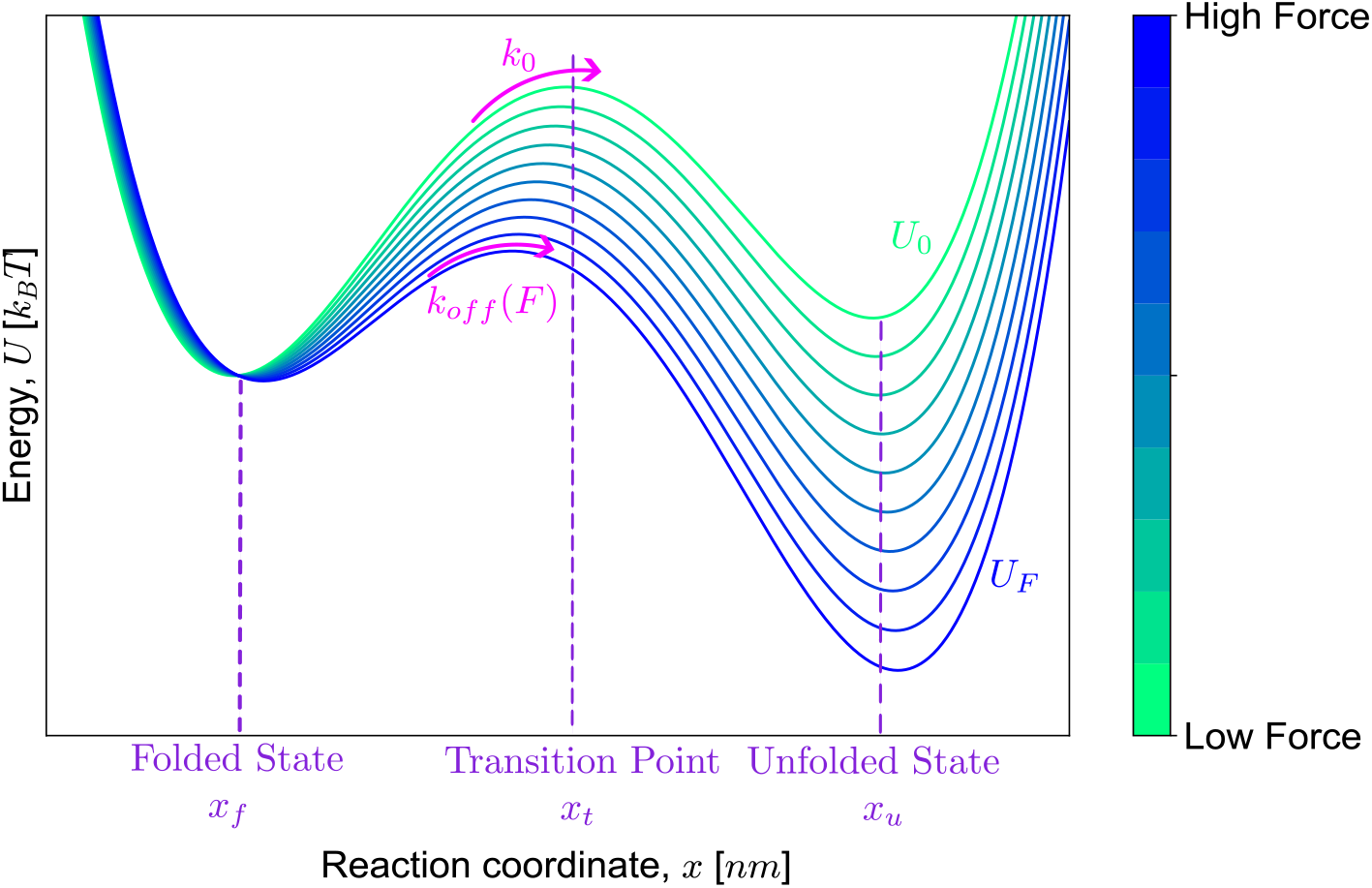
Schematic energy landscape along the reaction coordinate, where the intrinsic energy landscape *U*_0_ is titled by −*Fx* as the external force *F* increases, illustrated in blue.

An external force *F* results in a tilted energy landscape as shown in Fig. 2. The modified energy landscape is given by *U*_*F*_ (*x*) = *U*_0_(*x*) − *Fx* with a smaller barrier height. Additionally, the associated transition rate is denoted as *k*_*off*_ (*F*) and is related to the force-dependent lifetime *τ* (*F*), which is the time it takes to unfold under such force, through *k*_*off*_ (*F*) := 1*/*⟨*τ* (*F*)⟩; where ⟨·⟩ denotes the mean.

### The Dudko-Hummer Szabo model

In the context of twostate process, the transition rate *k*_*off*_ (*F*) can be obtained from the probability distribution *p*(*F*) of unfolding forces measured from constant speed mode as (16, 23),

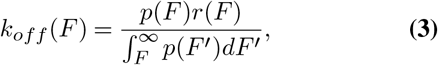

where 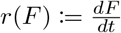 is the loading rate. This equation illustrates how the distributions of unfolding forces, measured under constant speed mode, can be converted into lifetime statistics measured in constant force mode, and is applicable regardless of the specific shape of the energy landscape.

The histogram of unfolding forces is constructed with equal bin width Δ*F*; starting from *F*_0_ and ending at *F*_*N*_ = *F*_0_ + *N* Δ*F*. Let the height of *k*th bin be,

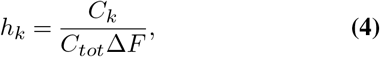

where *C*_*k*_ is the number of counts in *k*th bin and *C*_*tot*_ = ∑_*i*_ *C*_*i*_ is the total number of counts in the histogram. The transition rate *k*_*off*_ corresponding to the *k*th bin is found by (16),

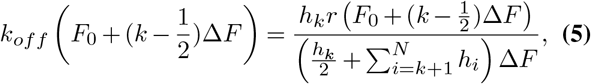

where 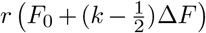 is the average loading rate of *k*th bin.

The Dudko-Hummer-Szabo (DHS) model (16) considers two different shapes of energy landscape to derive analytical expressions for *k*_*off*_ (*F*) using Kramers theory (22). The two energy landscapes are: 1) the cusp-like energy landscape which is described by 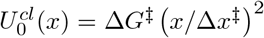 for *x < x*_*t*_ and 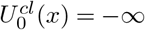 otherwise (24), and 2) the linear-cubic energy landscape which is described as 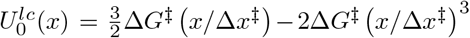 (23). The transition rate is given by,

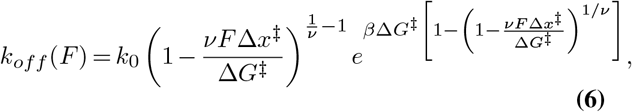

with ν = 1*/*2 or 2*/*3 for the cusp-like and linear-cubic energy landscape, respectively.

### Monte Carlo simulation

For the simulations, a protein of *N* folded domains is considered with one end attached to the substrate and the other end attached to the force probe. In the case of constant speed simulation, the base of the cantilever probe is moved away at a constant speed *v*; here the position of base of the cantilever *z* is initialized at zero and is updated every Δ*t* seconds. The protein extension *q* is determined by solving the equation,

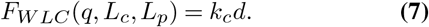

Here, *F*_*W LC*_(*q, L*_*c*_, *L*_*p*_) is the force applied on protein determined by the WLC model in Eq. 2. The transition rate *k*_*off*_ (*F*) is computed using Eq. 6, and the probability of a domain unfolding during the time interval Δ*t* is found by

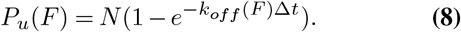

For determining unfolding events, a random number *η* is generated uniformly from 0 to 1 and is compared to the unfolding probability *P*_*u*_(*F*). No unfolding event is triggered if the random number is larger than *P*_*u*_(*F*) and the simulation will continue to the next time slot by adding time interval Δ*t*. Otherwise, one of the domains is unfolded, leading to a decrease in the number of folded domains, *N*, by 1, and the simulation continues to the next unfolding event if folded domains still remain. After each unfolding event, contour length and persistence length are updated by adding increments Δ*L*_*c*_, Δ*L*_*p*_ respectively. The constant speed pulling simulation algorithm is described in Algorithm 1, when performed with constant force, the force *F* is required instead of the speed *v*, and lines 2-4 in Algorithm 1 are skipped.

#### Algorithm 1 Protein Unfolding Monte Carlo Simulation

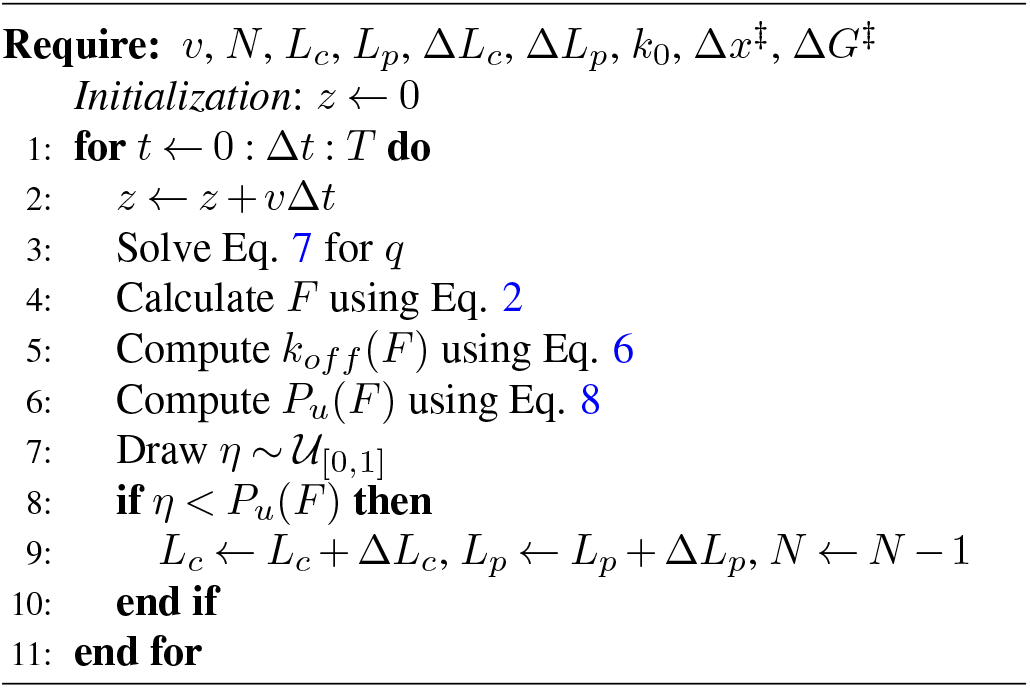

### Transition rates correction

The typical force feedback response time *τ*_*I*_ for Atomic Force Microscopy (AFM) is around 2 *ms* (25). Consequently, corrections are required to account for unfolding events that might not be observed due to the finiteness of the resolution (26). Under ideal conditions, with *τ*_*I*_ = 0, the transition rate is defined as the inverse of the average lifetime and is determined by,

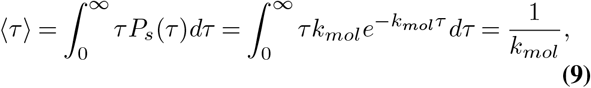

where the probability density of lifetime is given by 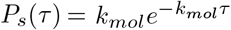 (according to Bell model (27)) and *k*_*mol*_ is the actual molecular transition rate. However, our observations of lifetimes are limited to *τ > τ*_*I*_. This limitation leads to the experimentally observed transition rate, *k*_*obs*_, which is determined by

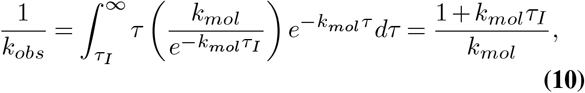

where the term enclosed within parentheses represents the normalization of the distribution over the restricted domain. By rearranging Eq. 10, we derive the correction formula that compensates for unobserved unfolding events attributed to the constraints posed by the finite instrumental response:

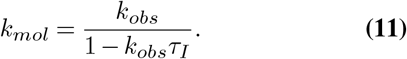

### Calibration protein

Titin I27O (Athena Enzyme Systems™), an AFM reference protein composed of eight repeats of the Ig 27 domain of human titin, serves as the calibration protein. The mode of unfolding forces at a pulling speed of 1000 *nm/s* is around 220 *pN* (Fig. S2c), closely matching the reported value of 225 *pN* (28). Additionally, energy landscape parameters align with values previously reported in the literature (23), where the literature values are represented by dashed lines in Fig. S2f-g.

## Results

### Constant speed and constant force experiments

AFM operational modes of constant speed and constant force were utilized for SMFS experiments. In experiments of a single protein molecule, one portion of the protein randomly attaches to the substrate and the rest of the molecule is left free to interact with the force probe, as illustrated by state 1 in Fig. 3a. The cantilever to substrate distance, *z*, is retracted at a constant rate in constant speed mode or is controlled to maintain the cantilever deflection *d* constant, thus ensuring a constant force on protein, in constant force mode. Under such applied mechanical tension, the protein is stretched, transitioning into state 2 in Fig. 3a. Subsequently, one of the folded protein domains unfolds stochastically, leading to state 3, and the cantilever continues to stretch the protein to state 4 in Fig. 3a. This process is repeated until all domains are unfolded or the connection between the cantilever and the substrate is broken.

**Fig. 3.**
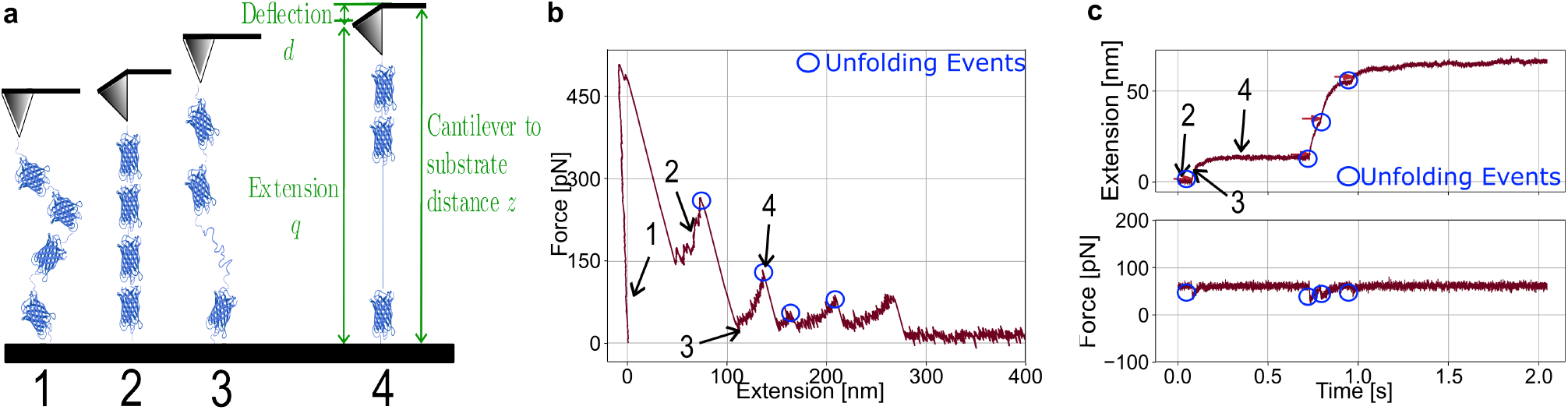
SMFS using two operational modes of AFM, constant speed and constant force. (a) Cartoon representation of the unfolding experiment: a molecule is attached between the substrate and the cantilever tip, experiencing stretching through states 1-4. (b) A representative force-extension curve of molecule insect UtrN-R3 from constant speed mode. (c) A representative curve of molecule insect UtrN-R3 from constant force mode, with the extension-time curve in the top panel and the force-time curve in the bottom panel. Unfolding events are highlighted in blue circles and the corresponding states are indicated by arrows in both (b) and (c).

In constant speed mode, the applied force drops abruptly when a domain unfolds, resulting in a saw-tooth pattern in the force versus extension curves. Fig. 3b illustrates a representative force-extension curve with unfolding events highlighted in blue circles. The states indicated by arrows are aligned with the cartoon schematic in Fig. 3a. In constant force mode, unfolding events are identified by discrete steps in the molecular extension accompanied by corresponding spikes in force, as illustrated in Fig. 3c. These unfolding events are marked with blue circles and a staircase-like pattern can be observed in the extension-time curve.

### Relating two operational modes via the DHS model

In constant speed mode, unfolding events were characterized by unfolding forces, leading to statistics of force magnitude data as shown in Fig. 4b. In the current study, data were collected under different pulling speeds (500, 1000, 2000, and 5000 *nm/s*) with lines representing the kernel density estimation of the corresponding histograms. Here, a Gaussian kernel was employed to estimate the underlying probability distribution of the data. While in constant force mode, the parameter of interest is the lifetime of a bound state which is the time for an unfolding event to occur from the moment the protein is subject to tension. In Fig. 4d, individual lifetimes of forces from 50 to 90 *pN* are represented as colored dots, with the brown line indicating the average lifetime.

**Fig. 4.**
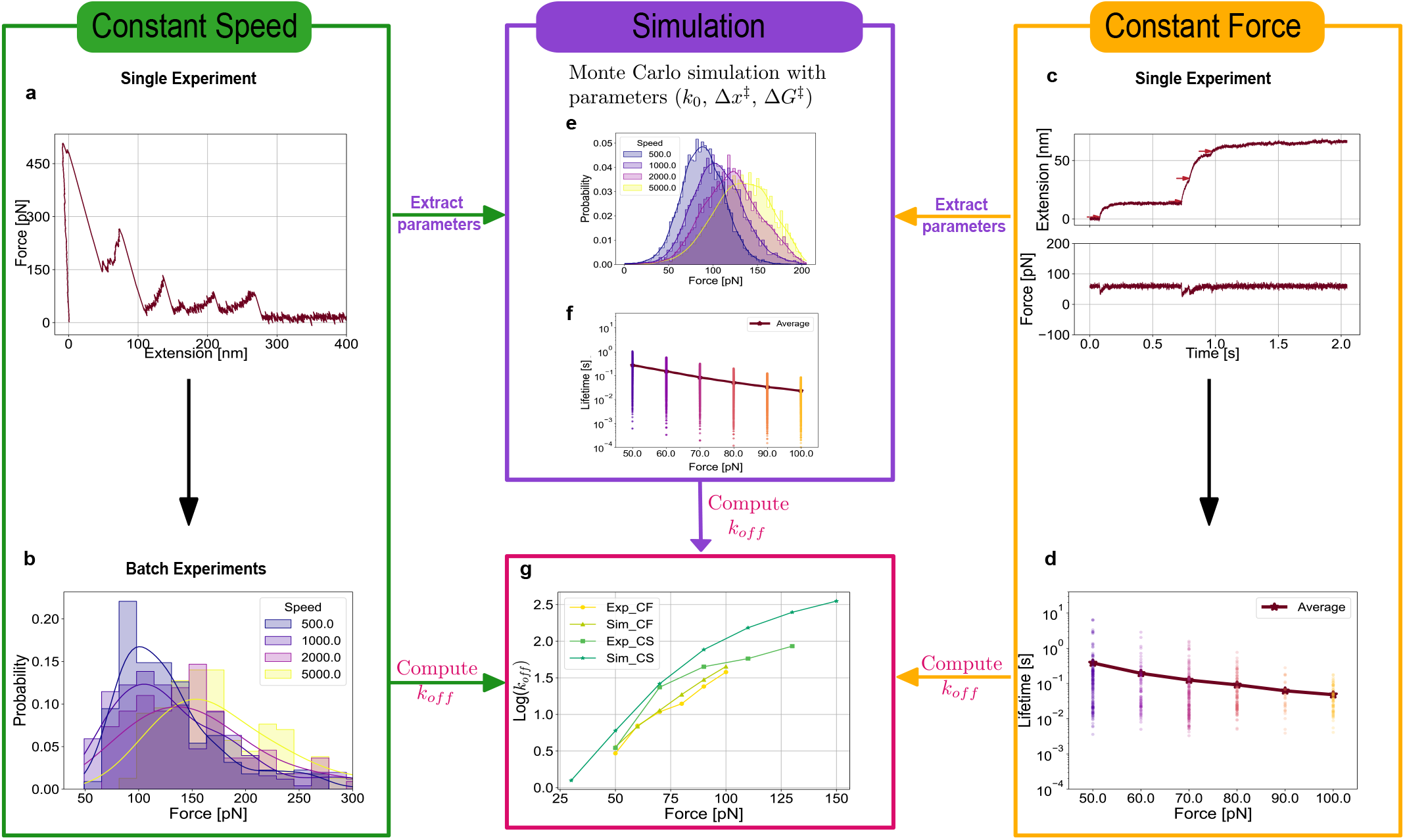
Consistent results using constant speed and constant force modes for insect UtrN-R3. (a) A representative force-extension curve from constant speed mode. (b) Unfolding force statistics of different pulling speeds ranging from 500 to 5000 *nm/s* with lines representing the kernel density estimation of the corresponding histograms. (c) A representative extension-time curve and force-time curve from constant force mode. (d) Measured lifetime from all unfolding events (colored dots) with the brown line representing the lifetime average, at forces changing from 50 to 100 *pN*. (e) Unfolding forces histograms along with kernel density estimations of simulation data at speeds ranging from 500 to 5000 *nm/s*. (f) Lifetime statistics of simulation data at forces changing from 50 to 100 *pN*. Individual lifetimes are represented by colored dots with the average shown as the brown line. (g) Transition rates *k*_*off*_ (*F*) for both experimental and simulated data using constant force and constant speed modes.

The DHS model was utilized to relate obtained statistics from the two different operational modes. Initially, the unfolding force distributions from different pulling speeds were used to estimate transition rate *k*_*off*_ (*F*) using Eq. 5 (See Fig. S6c). The average transition rate across all pulling speeds is presented in Fig. 4g, represented by the green square line. Subsequently, transition rates were determined from the lifetime statistics at different regulated forces in a constant force mode using Eq. 9 followed by the correction with Eq. 11. This is illustrated by the yellow line in Fig. 4g. Resulting transition rates were closely aligned with each other. To quantify this consistency, we used the coefficient of variation (CV), defined as the ratio of the standard deviation to the mean. Here the largest CV for *log*(*k*_*off*_) is 0.26 at 50 *pN*.

### Validating experimental data with Monte Carlo simulations

Our experimental data show consistent results using constant speed and constant force modes. To further validate our results, Monte Carlo simulations were conducted. However, these simulations require molecular parameters like contour length, persistence length, and their corresponding increments, which are estimated from experiments and molecule structure. Additionally, energy landscape parameters, including *k*_0_, Δ*G*^‡^, and Δ*x*^‡^, are essential and were extracted from curve fitting of Eq. 6 using the least squares method.

For the DHS model, average of experimental fitted parameters of the cusp-like energy landscape shape (*v* = 1*/*2) reported in Table 1 were utilized. Monte Carlo simulations in both constant speed and constant force modes as detailed in Algorithm 1 were conducted. The simulated force magnitude statistics and corresponding kernel density estimation are illustrated in Fig.4e. In Fig.4f. Subsequently, the transition rates were estimated using Monte Carlo simulations (Fig. 4g). Here the green line with triangles represents the average transition rate across different pulling speeds, with detailed transition rates from various pulling speeds in Fig. S6d. Notably, the transition rates were closely aligned (with CV*<* 0.3 in log(*k*_*off*_)) across four different scenarios: experimental constant speed, experimental constant force, simulated constant speed, and simulated constant force, as evidenced by Fig. 4g. This entire procedure is calibrated using Titin I27O (Fig. S2).

**Table 1.** DHS model parameters of three molecules: insect UtrN-R3, bacterial UtrN-R3, and bacterial DysN-R3.

| Molecules | DHS $\nu = 1/2$ | | | DHS $\nu = 2/3$ | | |
| --- | --- | --- | --- | --- | --- | --- |
| | $\ln(k_0)$ | $\Delta x^\ddagger$ [nm] | $\Delta G^\ddagger$ [$k_B T$] | $\ln(k_0)$ | $\Delta G^\ddagger$ [$k_B T$] | $\Delta G^\ddagger$ [$k_B T$] |
| Insect UtrN-R3 | −2.501 | 0.375 | 9.300 | −2.250 | 0.330 | 8.100 |
| Bacterial UtrN-R3 | −7.040 | 0.850 | 14.300 | −5.991 | 0.700 | 13.000 |
| Bacterial DysN-R3 | −4.501 | 0.600 | 11.500 | −3.450 | 0.480 | 10.000 |

<sup>a</sup> Energy landscapes depicted in Fig. 6 are based on the parameters provided here.
<sup>b</sup> These parameters are used in Monte Carlo simulations.
<sup>c</sup> They represent the average parameters obtained from experimental constant speed and constant force data, reported in S2-S4.

### Biological repeats reveal consistency among four scenarios

A single biological repeat of insect UtrN-R3 was initially utilized, and subsequent experiments and simulations of insect UtrN-R3 were expanded to include 3 biological repeats to increase the consistency of transition rates across various scenarios: experimental constant speed, experimental constant force, simulated constant speed, and simulated constant force. Simulations were conducted independently with parameters listed in Table 1. The individual lifetimes are represented by colored dots, with each color corresponding to the same biological repeat, and the lifetime averages are indicated by red triangles, calculated from identical forces and repeats. The lifetime averages exhibit a high degree of consistency among different biological repeats under the same force, in both experimental data (Fig. 5e) and simulation data (Fig. 5f). The largest CV observed for experimental data is 0.08 at 60 *pN*, while for simulation data it is 0.004 at 50 *pN*. Experimental lifetimes are truncated below approximately 0.01 *s*, while the simulated lifetimes are not. This discrepancy is attributed to the finite instrumental response, causing experimental lifetime averages to be slightly larger than simulated averages; Eq. 11 was employed to correct such bias. Unfolding force statistics are presented using violin plots for both experimental (Fig. 5g) and simulation data (Fig. 5h). The violins illustrate data distributions using kernel density estimation as black lines on each side, with the width of each curve indicating the relative frequency of data points. Meanwhile, black stars signify the positions of values with the highest probability, visualized by the widest section of the violin plot. The distributions are very similar to each other under the same pulling speed across different biological repeats for both experimental (CV*<* 0.08) and simulated data (CV*<* 0.02), with simulation data displaying fewer differences. The most probable values (black stars) are consistent between experiment and simulation. For example, at a pulling speed of 2000 *nm/s*, the unfolding forces are 123.7 ± 9.4 *pN* and 119.6 ± 2.4 *pN* for experimental and simulated data, respectively. However, a tail of high magnitudes is evident from experimental data, which may result from factors such as multi-molecule effects, variations in homogeneity, or bond interactions.

**Fig. 5.**
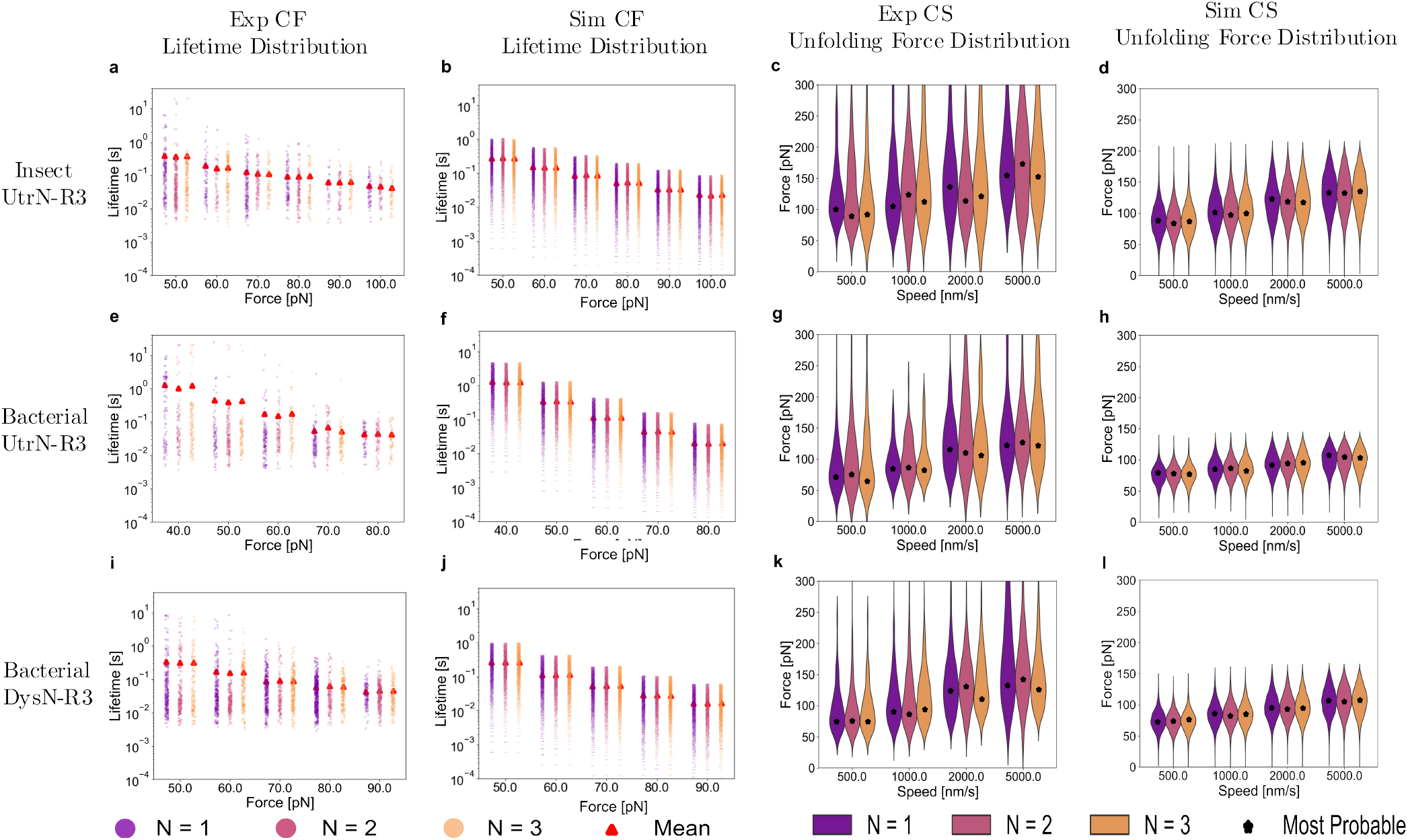
A summary of statistics of *N* = 3 biological repeats from four distinct scenarios, organized into columns representing experimental constant force, simulated constant force, experimental constant speed, and simulated constant speed from left to right. Each row presents results for different molecules, namely insect UtrN-R3, bacterial UtrN-R3, and bacterial DysN-R3 from top to bottom. The data from three biological repeats are visualized in purple, pink, and yellow. Individual lifetimes are represented by colored dots with red triangles indicating the lifetime average. Unfolding force distributions are depicted using violin plots, where black stars signify the positions of values with the highest probability. The violins illustrate data distributions using kernel density estimation as black lines on each side, with the width of each curve indicating the relative frequency of data points.

The transition rates *k*_*off*_ were computed using data from different biological repeats, and the mean values of *k*_*off*_ across these repeats are illustrated in Fig. S5d, with error bars representing standard deviations. Corresponding energy landscapes, fitted with both cusp-like and linear cubic energy landscape shapes, are depicted in Fig. S5e-f. The shaded areas in these plots indicate standard deviations of extracted parameters observed across various biological repeats, with associated parameters reported in SI Table S2. The above analysis and data on insect UtrN-R3 data demonstrate consistent results across four distinct scenarios. We then extend our investigation to two additional molecules: bacterial UtrN-R3, and bacterial DysN-R3.

### Mechanical stiffness of bacterial UtrN-R3 is significantly lower in comparison to insect UtrN-R3 while remaining consistent across four scenarios

Our earlier research (13) demonstrated that the unfolding forces of UtrNR3 are influenced by expression systems. In particular, insect UtrN-R3 exhibited higher unfolding forces compared to bacterial UtrN-R3 when subjected to a constant pulling speed of 1000 *nm/s*. Building upon the data obtained at 1000 *nm/s* from our previous study, we conducted additional experiments at pulling speeds of 500, 1000, 2000, and 5000 *nm/s* for both insect and bacterial UtrN-R3. The distributions of unfolding forces for bacterial UtrN-R3 shifted towards higher values at higher pulling speeds, as illustrated in Fig. 5g. Notably, the most probable unfolding forces consistently remained 25.5 ± 10.9 *pN* less than those observed in insect UtrN-R3 under equivalent speeds. Typical curves for bacterial UtrN-R3 are available in SI Fig. S3g-i for constant speed mode and SI Fig. S4g-i for constant force mode. Since bacterial UtrN-R3 exhibited lower unfolding forces as compared to insect UtrN-R3, correspondingly lower forces were applied to bacterial UtrN-R3 in constant force mode, with lifetime statistics in Fig. 5e. The lifetime averages were 0.038 ± 0.024 *s* smaller compared to insect UtrN-R3 under the same force. Unfolding forces (CV *<* 0.06) and lifetime statistics (CV *<* 0.13) were consistent across biological replicates. Parameters for *v* = 1*/*2 reported in Table 1 were taken to run Monte Carlo simulations. The results, including transition rates (Fig. S5d), fitted energy landscapes (Fig. S5e-f) and corresponding parameters (SI Table S3) demonstrate consistency among different scenarios.

We provide a comprehensive overview of transition rates *k*_*off*_ in Fig. 6a. Simultaneously, corresponding energy landscapes with both cusp-like and linear cubic shapes are plotted in Fig. 6b-c, with parameters in Table 1. To quantify the steepness of the energy landscape, parameters from the cusp-like shape (ν = 1*/*2 in Table 1) were considered. Assuming an external force of 20 *pN* is applied to tilt the energy landscape, the tilted energy barrier of insect UtrNR3 becomes approximately 1.8 *k*_*B*_*T* (using the relationship *U*_*F*_ (*x*) = *U*_0_(*x*) − *Fx*), while the energy barrier of bacterial UtrN-R3 changes to approximately −2.7 *k*_*B*_*T*. Since the energy barrier decreases more slowly under external forces in insect UtrN-R3, despite the energy barrier Δ*G*^‡^ of insect UtrN-R3 being 2.2 *k*_*B*_*T* smaller than bacterial UtrN-R3 when no external force is applied, insect UtrN-R3 is mechanically stiffer than bacterial UtrN-R3.

**Fig. 6.**
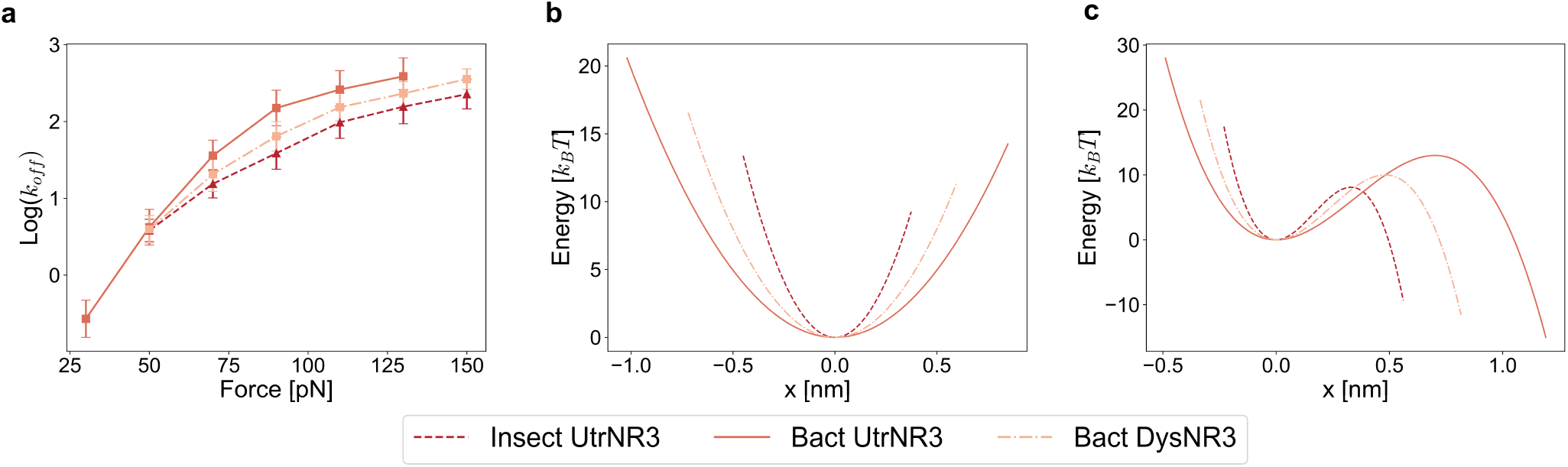
A summary of results across all three molecules, insect UtrN-R3, bacterial UtrN-R3, and bacterial DysN-R3. (a) *k*_*off*_-vs-*F* curves with mean and standard deviations across different scenarios and biological replicates denoted by lines and error bars, respectively. Fitted energy landscapes with (b) ν = 1*/*2 and (c) ν = 2*/*3.

We conducted the Mann-Whitney U test (29), which tests the null hypothesis that two samples share the same distribution, on the constant speed data with the level of confidence *α* = 0.05. Across four different speeds, all p-values comparing insect and bacterial UtrN-R3 are below 0.0001(Fig. 7a, Table S5), indicating significantly different unfolding force distributions between these two molecules. For constant force data, the t-test (30), which is a test for the null hypothesis that 2 independent samples have identical mean values, is employed with *α* = 0.05. All p-values comparing insect and bacterial UtrN-R3 under forces from 50 to 80 *pN* are all smaller than 0.001 (Fig. 7b, Table S5). This suggests distinct lifetime averages between these two molecules. Taken together, we can conclude that insect and bacterial UtrN-R3 are significantly different than each other in terms of mechanical properties.

**Fig. 7.**
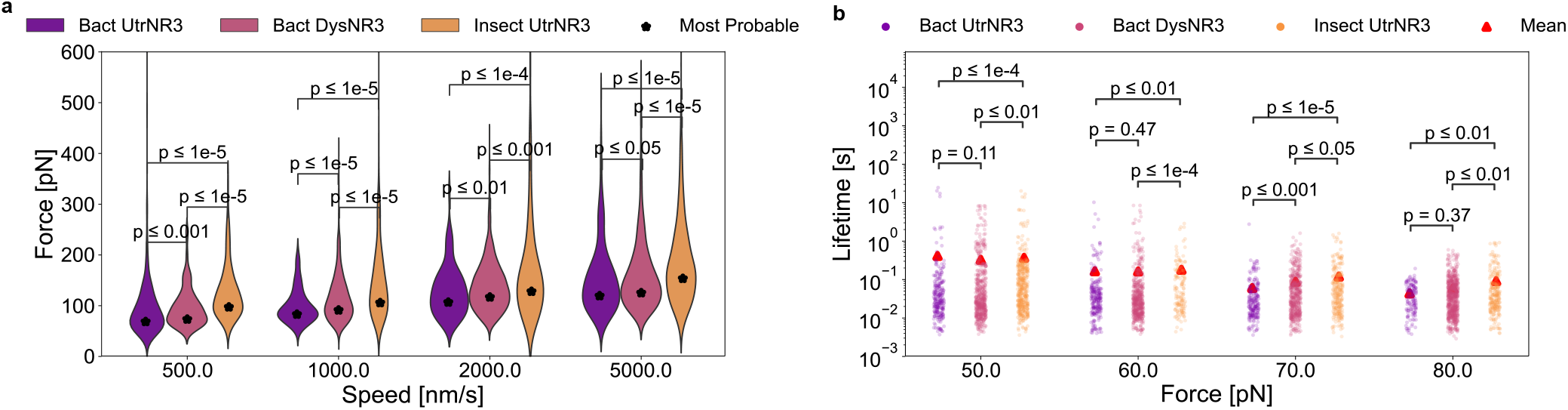
Statistics for all three molecules, bacterial UtrN-R3, bacterial DysN-R3, and insect UtrN-R3, from both (a) constant speed, and (b) constant force data. The p-values over brackets correspond to comparisons of paired data.

### Stiffness of bacterial DysN-R3 is significantly lower than those of insect UtrN-R3, but exhibits similarity to bacterial UtrN-R3

We have previously reported that bacterial DysR8-15 exhibits comparable values to bacterial UtrNR3 with unfolding forces of 100.1 ± 7.0 *pN* when consistently pulled at 1000 *nm/s*. Here, bacterial DysN-R3 was utilized to make a direct comparison with UtrN-R3 expressed in both insect and bacteria cells. Experimental investigations were conducted in both constant speed and constant force modes, followed by simulations after extracting energy landscape parameters (Fig. 5 i-l). The unfolding forces and lifetime statistics remained consistent (CV*<* 0.1) across different biological replicates and scenarios. Moreover, the results demonstrate consistency once again in terms of transition rate, with CV*<* 0.3 in *log*(*k*_*off*_) (Fig. S5g), and fitted energy parameters (SI Table S4), along with corresponding energy landscapes (Fig. 3h-i).

Under experimental constant speed mode at 1000 *nm/s*, the most probable value of unfolding forces across 3 biological replicates for bacterial DysN-R3 is reported to be 90.3 ± 3.2 *pN* (Fig. 5k). In contrast, the most probable values for insect and bacterial UtrN-R3 are 113.7 ±7.6 *pN* and 84.5 ±1.6 *pN*, respectively. The order of magnitude of unfolding forces holds under various speeds, with bacterial UtrN-R3, bacterial DysN-R3, and insect UtrN-R3 showing a trend from smallest to largest (Fig. 7a). Similarly, the same order holds in terms of lifetime averages(Fig. 7b). For example, the lifetime averages are 0.043 ± 0.001 *s*, 0.059 ± 0.002 *s*, and 0.092 ± 0.001 *s* for bacterial UtrN-R3, bacterial DysN-R3, and insect UtrNR3, respectively. Additionally, both cusp-like and linear cubic energy landscapes of bacterial DysN-R3 (dashdot lines in Fig. 6b-c) are located between those of insect and bacterial UtrN-R3. Therefore, we can conclude that bacterial DysNR3 stiffness is between insect and bacterial UtrN-R3.

To quantify the difference among the three molecules, we conducted the Mann-Whitney U test with constant speed data on two pairs: bacterial UtrN-R3 and bacterial DysN-R3, and bacterial DysN-R3 and insect UtrN-R3. Despite the p-values of both pairs being less than 0.05 under each speed, the pvalues comparing bacterial UtrN-R3 and bacterial DysN-R3 are consistently lower than those comparing bacterial DysNR3 and insect UtrN-R3 (Fig. 7a, Table S5). Similarly, the t-test was used for the constant force data. All the p-values of the pair bacterial DysN-R3 and insect UtrN-R3 are less than 0.05, indicating differences between these two proteins (Fig. 7b, Table S5). However, the p-values between bacterial UtrN-R3 and bacterial DysN-R3 are larger than 0.05 at forces of 50 and 60 *pN*. In summary, bacterial DysNR3 significantly differs from insect UtrN-R3 in every speed and force, but bacterial DysN-R3 is not significantly different from bacterial UtrN-R3 at some forces. Additionally, the p-values between bacterial DysN-R3 and bacterial UtrN-R3 are higher compared to those between bacterial DysN-R3 and insect UtrN-R3. This suggests that the mechanical properties of bacterial DysN-R3 are more similar to bacterial UtrN-R3 while lying between bacterial and insect UtrN-R3.

## Discussion

In our experiments, we utilized both constant speed and constant force modes. The force magnitude at which the domain unfolds and the lifetime, the time of unfolding, are recorded in constant speed and constant force modes, respectively. All experiments were carried out with *N* = 3 biological repeats, showcasing consistent unfolding force and lifetime statistics among different replicates under the same conditions (Fig. 5). Our data from different modes exhibit consistency in transition rates *k*_*off*_ using the DHS model, which is further validated with Monte Carlo simulations (Fig. S5).

We previously reported significantly different constant speed unfolding force statistics for insect versus bacterial UtrNR3, and showed that phosphorylation impacts the mechanical stiffness of UtrN-R3 (13). Specifically, phosphorylated insect UtrN-R3 exhibits higher unfolding forces compared to unphosphorylated bacterial UtrN-R3. However, these findings were limited to data collected at 1000 *nm/s* in constant speed mode. Here we conducted additional experiments involving both insect and bacterial UtrN-R3 under various speeds in constant speed mode, as well as under different forces in constant force mode. Our data confirms that insect UtrN-R3 exhibits significantly stiffer mechanical properties than bacterial UtrN-R3. The most probable unfolding forces of insect UtrN-R3 are 25.5 ± 10.9 *pN* higher than those of bacterial UtrN-R3 under the same pulling speeds, and the lifetime averages are 0.038 ±0.024 s longer (Fig. 5). Energy landscapes confirm this result (Fig. 6k-l), with insect UtrN-R3 having a steeper energy barrier. Original to this study, we demonstrate that the mechanical properties of bacterial DysN-R3 lie between insect and bacterial UtrN-R3. This is supported by unfolding force and lifetime statistics (Fig. 5), as well as the extracted energy landscapes (Fig. 6). Furthermore, bacterial DysN-R3 exhibits similarities to bacterial UtrN-R3 rather than insect UtrN-R3, as evidenced by the p-values in Fig. 7 and Table S5.

In conclusion, our study quantifies the mechanical properties of insect UtrN-R3, bacterial UtrN-R3, and bacterial DysNR3 using both constant speed and constant force modes. Data from different modes show consistency via the DHS model. Our data, including unfolding forces, lifetimes, and energy landscape parameters, suggest that when compared directly, homologous fragments of dystrophin and utrophin are more similar in mechanical properties than predicted from prior studies across laboratories. Our data suggest that from a mechanical perspective, utrophin is a good therapeutic surrogate for dystrophin deficiency in DMD.

## Author Contributions

C.H., J.M.M., J.M.E., and M.V.S. designed the research. C.H., R.A.S., and J.V. conducted experiments. C.H. analyzed the data. C.H. and M.V.S. wrote the article. All authors participated in reviewing/editing of the manuscript and approved the final draft.

## Acknowledgments

This project was supported by funding from NIH (R01AR042423 and 5T32AR007612) and the Muscular Dystrophy Association (628925).

