## Supplemental Files for "Two operational modes of atomic force microscopy reveal similar mechanical properties for homologous regions of dystrophin and utrophin"

DRAFT

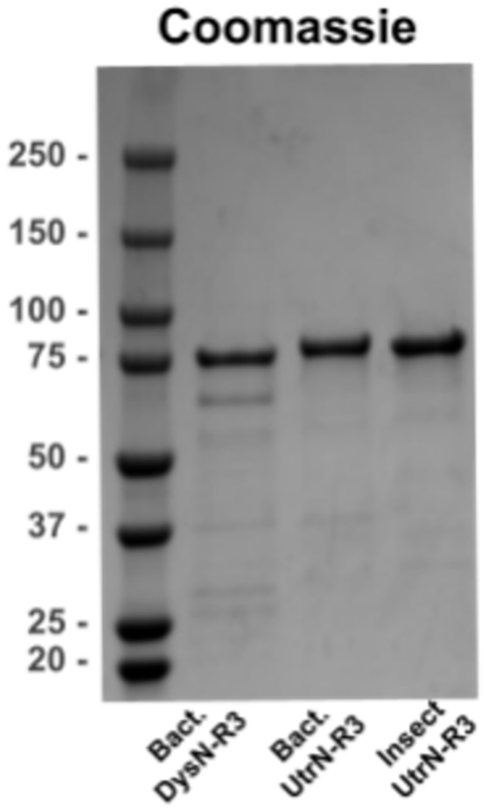


**Fig. 1.** Coomassie blue-stained SDS polyacrylamide gel with 2 $\mu g$ purified bacterial. DysN- R3, bacterial UtrN-R3, and insect UtrN-R3. Molecular weight standards $(\times 10^3)$ are shown on the left.

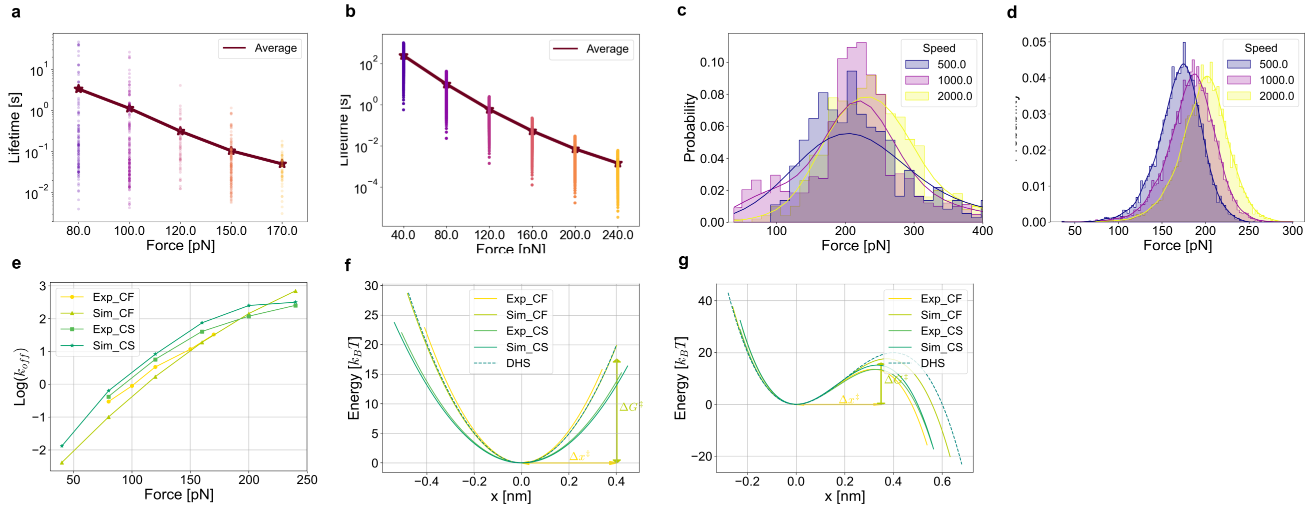


**Fig. 2.** Summary of Titin I27O characterization. (a) Measured experimental lifetime from all unfolding events (colored dots) with the brown line representing the lifetime average, at forces ranging from $80$ to $170\ pN$. Typical curves are provided in SI Fig. 4a-c. (b) Lifetime statistics of simulation data at forces changing from $40$ to $240\ pN$, with energy landscape parameters adopted from (1). (c) Unfolding force statistics of different pulling speeds ($500$, $1000$, and $2000\ nm/s$) with lines representing the kernel density estimation of the corresponding histograms. (d) Unfolding forces histograms along with kernel density estimations of simulation data at speeds including $500$, $1000$, and $2000\ nm/s$. Representative curves are displayed in SI Fig. 3a-c. (e) Transition rates $k_{off}(F)$ for both experimental and simulated data using constant force and constant speed modes. Fitted energy landscapes with $\nu = 1/2$ and $\nu = 2/3$ are shown in (f) and (g), respectively. Here, $\Delta x^{\ddagger}$ and $\Delta G^{\ddagger}$ represent the averages across four different scenarios, as documented in Table 1.

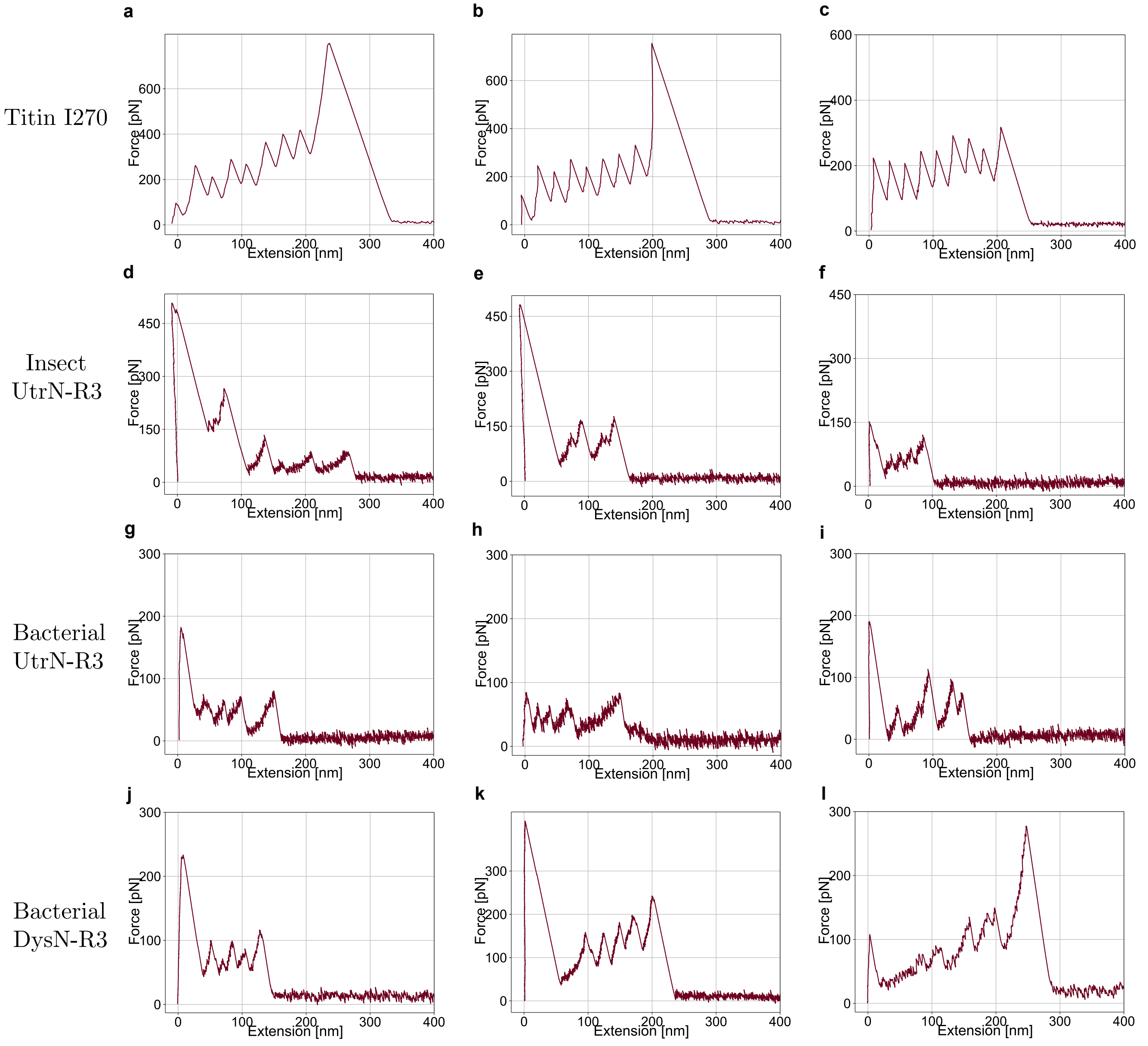


**Fig. 3.** Representative curves from constant speed experiments. Each row presents results for different molecules, including Titin I27O, insect UtrN-R3, bacterial UtrN-R3, and bacterial DysN-R3 from top to bottom.

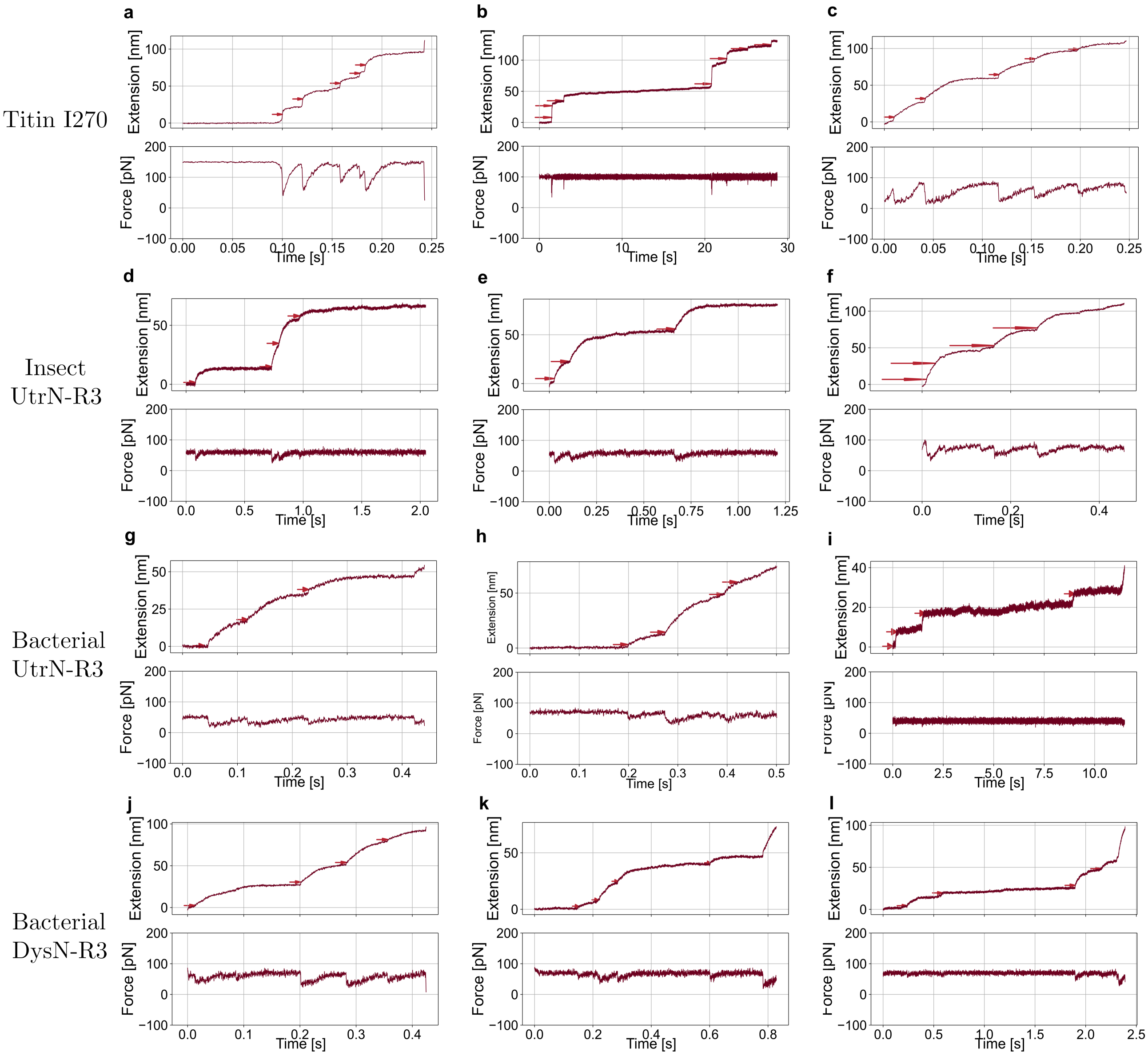


**Fig. 4.** Representative curves from constant force experiment. Each row represents results namely Titin I27O, insect UtrN-R3, bacterial UtrN-R3, and bacterial DysN-R3 from top to bottom.

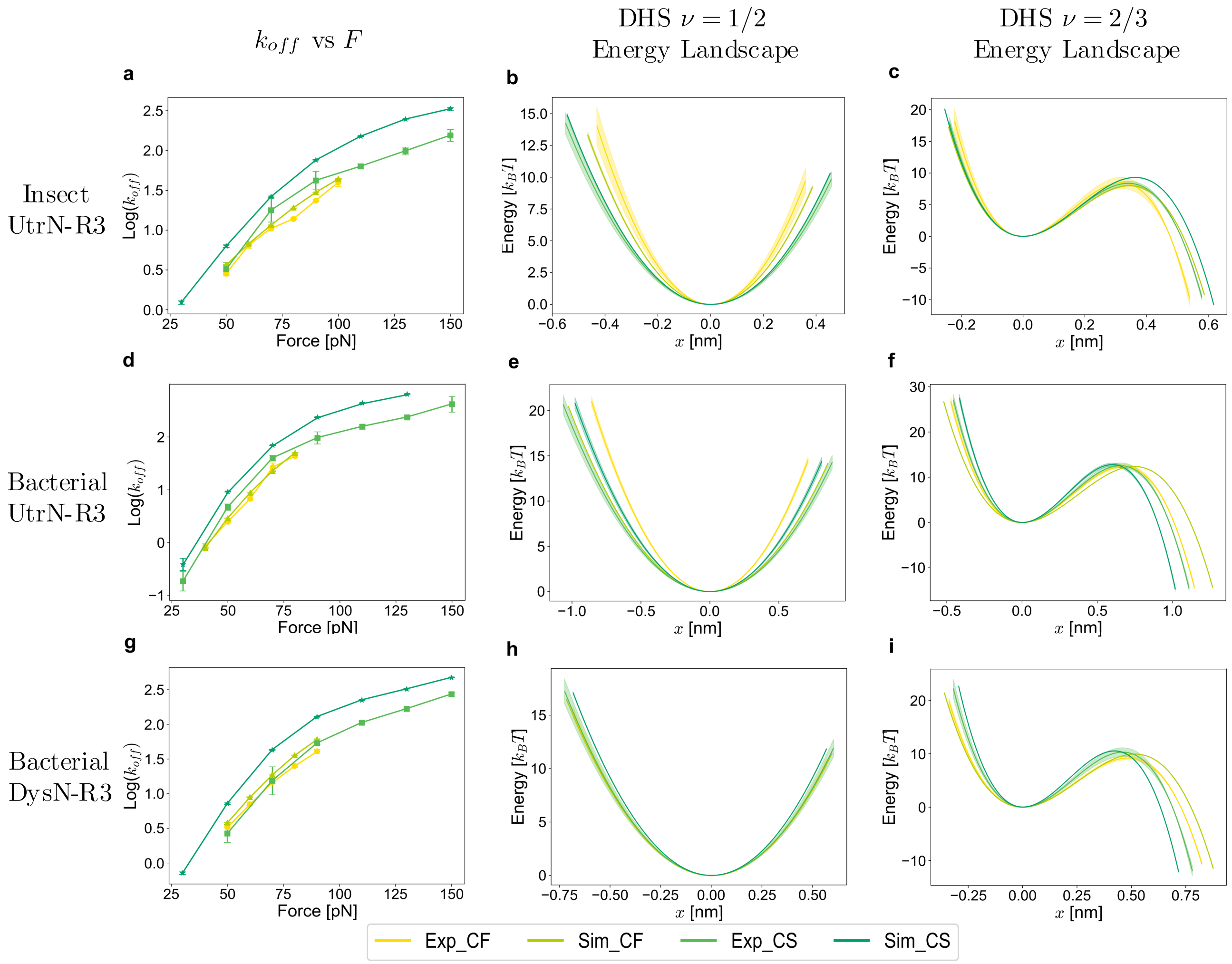


**Fig. 5.** A summary of results across four distinct scenarios: experimental constant speed, experimental constant force, simulated constant speed, and simulated constant force. Each row presents results for different molecules, namely insect UtrN-R3, bacterial UtrN-R3, and bacterial DysN-R3 from top to bottom. The graphs in the first column show $k_{off}$-vs-$F$ curves, while the second and third columns show fitted energy landscapes with $\nu = 1/2$ and $\nu = 2/3$, respectively.

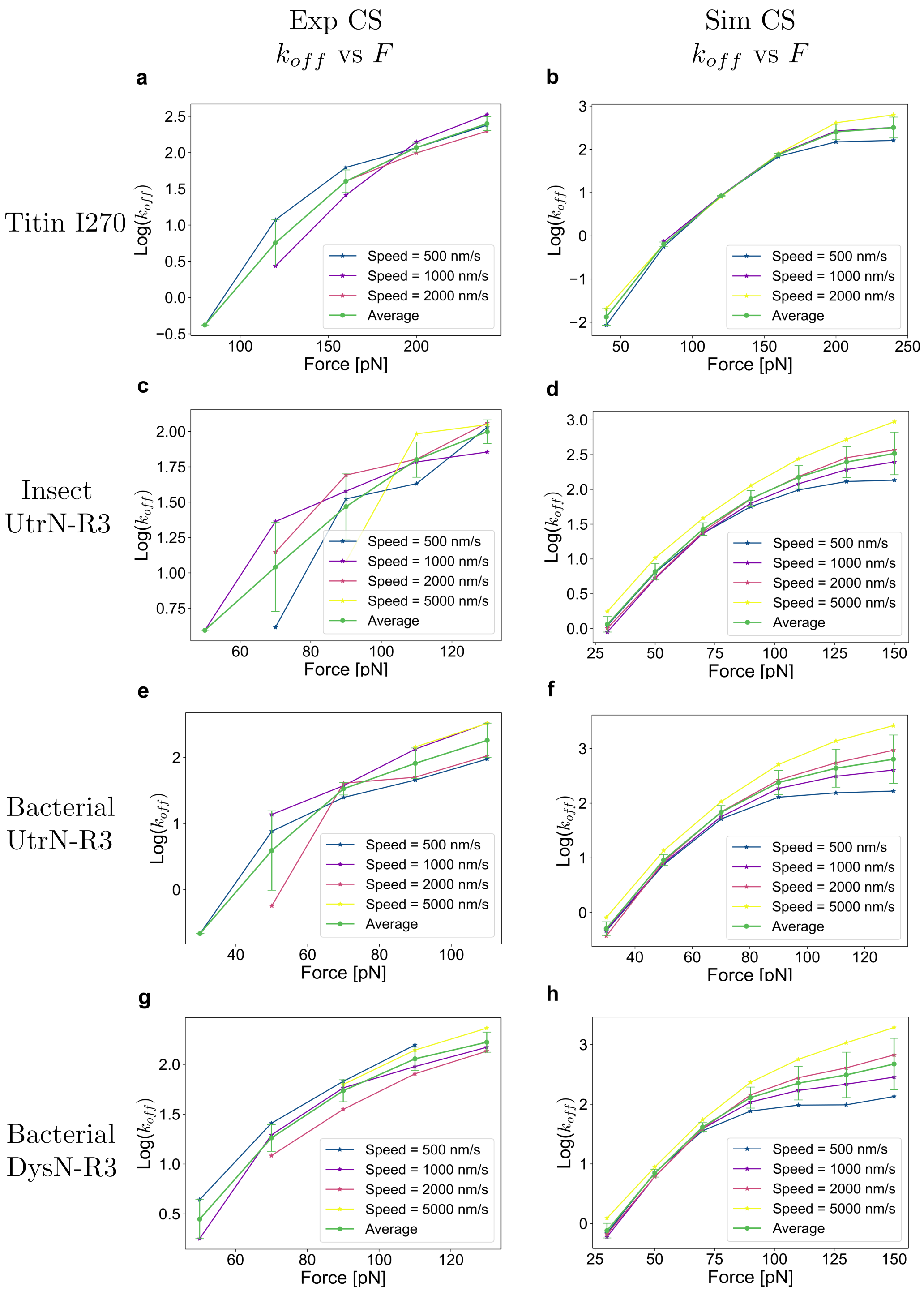


**Fig. 6.** Transition rates of constant speed mode from both experimental and simulated data for Titin I27, insect UtrN-R3, bacterial UtrN-R3, and bacterial DysN-R3. Here only the first biological repeat (N = 1) is reported, the other two repeats exhibit similar trends and are thus omitted. Each row presents results for different molecules, namely Titin I27O, insect UtrN-R3, bacterial UtrN-R3, and bacterial DysN-R3 from top to bottom. The first column utilizes experimental data while the second uses simulated data. The green line represents the average across different speeds, with the error bar indicating the standard deviation.

**Table 1.** Titin I27O modeling parameters

| Molecules | DHS $\nu = 1/2$ | | | DHS $\nu = 2/3$ | | |
|---|---|---|---|---|---|---|
| | $ln(k_0)$ | $\Delta x^{\ddagger}$ $[nm]$ | $\Delta G^{\ddagger}$ $[k_B T]$ | $ln(k_0)$ | $\Delta x^{\ddagger}$ $[nm]$ | $\Delta G^{\ddagger}$ $[k_B T]$ |
| Exp_CF | $-6.911$ | $0.342$ | $15.936$ | $-6.672$ | $0.317$ | $13.623$ |
| Sim_CF | $-9.090$ | $0.403$ | $19.970$ | $-8.852$ | $0.373$ | $17.586$ |
| Exp_CS | $-7.604$ | $0.423$ | $15.299$ | $-6.388$ | $0.329$ | $13.482$ |
| Sim_CS | $-8.385$ | $0.465$ | $17.01$ | $-6.954$ | $0.341$ | $15.506$ |

**Table 2.** Insect UtrN-R3 modeling parameters

| Molecules | DHS $\nu = 1/2$ | | | DHS $\nu = 2/3$ | | |
|---|---|---|---|---|---|---|
| | $ln(k_0)$ | $\Delta x^{\ddagger}$ $[nm]$ | $\Delta G^{\ddagger}$ $[k_B T]$ | $ln(k_0)$ | $\Delta x^{\ddagger}$ $[nm]$ | $\Delta G^{\ddagger}$ $[k_B T]$ |
| Exp_CF | $-2.445 \pm 0.301$ | $0.360 \pm 0.039$ | $9.758 \pm 0.301$ | $-2.185 \pm 0.241$ | $0.318 \pm 0.027$ | $8.472 \pm 0.882$ |
| Sim_CF | $-2.497 \pm 0.136$ | $0.388 \pm 0.018$ | $9.264 \pm 0.136$ | $-2.264 \pm 0.100$ | $0.345 \pm 0.012$ | $7.997 \pm 0.094$ |
| Exp_CS | $-3.083 \pm 0.605$ | $0.459 \pm 0.072$ | $9.901 \pm 0.588$ | $-2.142 \pm 0.151$ | $0.341 \pm 0.022$ | $8.355 \pm 0.371$ |
| Sim_CS | $-2.603 \pm 0.063$ | $0.454 \pm 0.005$ | $10.375 \pm 0.078$ | $-2.031 \pm 0.046$ | $0.363 \pm 0.003$ | $9.322 \pm 0.067$ |

[a] Values are mean $\pm$ standard deviations from $N = 3$ biological repeats

**Table 3.** Bacterial UtrN-R3 modeling parameters

| Molecules | DHS $\nu = 1/2$ | | | DHS $\nu = 2/3$ | | |
|---|---|---|---|---|---|---|
| | $ln(k_0)$ | $\Delta x^{\ddagger}$ $[nm]$ | $\Delta G^{\ddagger}$ $[k_B T]$ | $ln(k_0)$ | $\Delta x^{\ddagger}$ $[nm]$ | $\Delta G^{\ddagger}$ $[k_B T]$ |
| Exp_CF | $-6.012 \pm 0.585$ | $0.715 \pm 0.063$ | $14.600 \pm 0.349$ | $-5.894 \pm 1.845$ | $0.599 \pm 0.060$ | $12.425 \pm 0.362$ |
| Sim_CF | $-6.951 \pm 0.146$ | $0.856 \pm 0.022$ | $14.228 \pm 0.038$ | $-6.418 \pm 0.098$ | $0.744 \pm 0.013$ | $12.416 \pm 0.022$ |
| Exp_CS | $-6.923 \pm 0.395$ | $0.887 \pm 0.014$ | $14.366 \pm 0.745$ | $-5.532 \pm 0.219$ | $0.652 \pm 0.021$ | $12.587 \pm 0.532$ |
| Sim_CS | $-5.798 \pm 0.414$ | $0.814 \pm 0.034$ | $14.470 \pm 0.454$ | $-4.440 \pm 0.302$ | $0.598 \pm 0.018$ | $12.787 \pm 0.359$ |

[a] Values are mean $\pm$ standard deviations from $N = 3$ biological repeats

**Table 4.** Bacterial DysN-R3 modeling parameters

| Molecules | DHS $\nu = 1/2$ | | | DHS $\nu = 2/3$ | | |
|---|---|---|---|---|---|---|
| | $ln(k_0)$ | $\Delta x^{\ddagger}$ $[nm]$ | $\Delta G^{\ddagger}$ $[k_B T]$ | $ln(k_0)$ | $\Delta x^{\ddagger}$ $[nms]$ | $\Delta G^{\ddagger}$ $[k_B T]$ |
| Exp_CF | $-4.184 \pm 0.194$ | $0.570 \pm 0.016$ | $10.614 \pm 0.371$ | $-3.711 \pm 0.212$ | $0.485 \pm 0.016$ | $9.196 \pm 0.333$ |
| Sim_CF | $-4.357 \pm 0.075$ | $0.597 \pm 0.010$ | $11.469 \pm 0.012$ | $-3.904 \pm 0.042$ | $0.515 \pm 0.005$ | $9.953 \pm 0.009$ |
| Exp_CS | $-4.804 \pm 0.748$ | $0.604 \pm 0.036$ | $11.983 \pm 0.818$ | $-3.705 \pm 0.791$ | $0.458 \pm 0.037$ | $10.309 \pm 0.821$ |
| Sim_CS | $-3.693 \pm 0.053$ | $0.570 \pm 0.005$ | $11.891 \pm 0.063$ | $-2.677 \pm 0.040$ | $0.422 \pm 0.003$ | $10.509 \pm 0.053$ |

[a] Values are mean $\pm$ standard deviations from $N = 3$ biological repeats

**Table 5.** The p-values for comparison among three molecules: bacterial UtrN-R3, bacterial DysN-R3, and insect UtrN-R3.

| Conditions | | Bacterial UtrN-R3 vs Insect UtrN-R3 | Bacterial UtrN-R3 vs Bacterial DysN-R3 | Bacterial DysN-R3 vs Insect UtrN-R3 |
|---|---|---|---|---|
| Constant Speed [nm/s] | 500 | **** | *** | **** |
| | 1000 | **** | **** | **** |
| | 2000 | **** | ** | *** |
| | 5000 | **** | * | **** |
| Constant Force [pN] | 50 | **** | 0.06 | *** |
| | 60 | **** | 0.59 | **** |
| | 70 | **** | **** | ** |
| | 80 | *** | * | * |

[a] p-value annotation legend: *: $0.01 < p \leq 0.05$; **: $0.001 < p \leq 0.01$; ***: $0.0001 < p \leq 0.001$; and ****: $p \leq 0.0001$

1. Olga K. Dudko, Gerhard Hummer, and Attila Szabo. Intrinsic rates and activation free energies from single-molecule pulling experiments. *Physical review letters*, 96(10):108101, 2006.
